# Limb apparent motion perception: modification by tDCS, and clinically or experimentally altered bodily states

**DOI:** 10.1101/2021.03.06.433957

**Authors:** Gianluca Saetta, Jasmine T. Ho, Robin Bekrater-Bodmann, Peter Brugger, H. Chris Dijkerman, Bigna Lenggenhager

**Affiliations:** Department of Psychology, University of Zurich, Switzerland; Department of Experimental Psychology, Utrecht University, The Netherlands; Department of Cognitive and Clinical Neuroscience, Central Institute of Mental Health, Medical Faculty Mannheim, Heidelberg University, Mannheim, Germany; Department of Psychiatry, Psychotherapy and Psychosomatics, University Hospital of Psychiatry (PUK), Zurich, Switzerland; Neuropsychology Unit, Valens Rehabilitation Centre, Valens, Switzerland

**Author notes:** Corresponding author: Gianluca Saetta, Department of Psychology, University of Zurich, Cognitive Neuropsychology, Binzmühlerstrasse, 14, CH-8050 Zurich, Switzerland, **Email**. Requests for reprints to Gianluca Saetta.

**Keywords:** embodied cognition, apparent motion perception, tDCS, lower limb amputation, body integrity dysphoria

## Abstract

Limb apparent motion perception (LAMP) refers to the illusory visual perception of a moving limb upon observing two rapidly alternating photographs depicting the same limb in two different postures. Fast stimulus onset asynchronies (SOAs) induce the more visually guided perception of physically impossible movements. Slow SOAs induce the perception of physically possible movements. According to the motor theory of LAMP, the latter perception depends upon the observers’ sensorimotor representations. Here, we tested this theory in two independent studies by performing a central (study 1) and peripheral (study 2) manipulation of the body’s sensorimotor states during two LAMP tasks. In the first sham-controlled transcranial direct current stimulation between-subject designed study, we observed that the dampening of left sensorimotor cortex activity through cathodal stimulation biased LAMP towards the more visually guided perception of physically impossible movements for stimulus pairs at slow SOAs. In the second, online within-subject designed study, we tested three participant groups twice: (1) individuals with an acquired lower limb amputation, either while wearing or not wearing their prosthesis; (2) individuals with body integrity dysphoria (i.e., with a desire for amputation of a healthy leg) while sitting in a regular position or binding up the undesired leg (to simulate the desired amputation); (3) able-bodied individuals while sitting in a normal position or sitting on one of their legs. We found that the momentary sensorimotor state crucially impacted LAMP in all groups. Taken together, the results of these two studies substantiate the motor theory of LAMP.

## Introduction

Embodied cognition theory advocates an essential contribution of the human body’s structure, functionality, and sensorimotor state on perception, action, and cognition (Barsalou, 2010; Bechara & Damasio, 2005). In this framework, the repertoire of feasible movements and basic principles of physics, such as the implicit notion of mutual impenetrability of two solid entities (the law of impenetrability, Heinemann, 1945), may guide visual perception of body movements (Saetta et al., 2018). Since such movements are often partially occluded, accurate prediction about them is crucial (Kilner et al., 2007). Accordingly, there are dedicated mechanisms to extract the perception of coherent and dynamic bodily movement trajectories from partially occluded or even static visual cues (Downing et al., 2001; Giese & Poggio, 2003). For instance, human movement kinematics can be inferred from point-light displays applied to a human walker’s joints in an otherwise darkened space (Blake & Shiffrar, 2007a). Moreover, the presentation of static photographs implying motion (e.g., an actor jumping off a cliff) biases spatial memory about the direction of the implied motion (Kourtzi & Shiffrar, 1999; Verfaillie & Daems, 2002).

A compelling illustration of such predictive mechanisms is the so-called limb apparent motion perception (LAMP, Shiffrar & Freyd, 1990). LAMP refers to the illusory completion of an actor’s limb movements generated by alternating two motionless pictures depicting the same limb in two different positions. In LAMP tasks, the perception of the limb’s movement trajectory is manipulated by the time interval between the two pictures’ onset (i.e., stimulus-onset asynchrony, SOA). *Fast SOAs* typically induce the perception of a short movement trajectory that reflects physically impossible movements: In violation of the law of impenetrability, the limb is perceived as going trough a solid object, or, in violation of the biomechanical constraints, the limb is perceived to move along a short angle of rotation incompatible with the joint’s biomechanics. Therefore, this kind of perception has been supposed to be *visually guided*, i.e., relying on visual perception that is independent of the observer’s motor capabilities (Saetta et al., 2018; Vannuscorps & Caramazza, 2016; Shiffrar & Freyd, 1990).

*Slow SOAs,* on the other hand, induce the perception of a large movement trajectory that reflects physically possible movements (i.e., the limb is perceived as moving around the solid object or as rotating around a joint consistent with its biomechanical constraints). This perception is thought to be *sensorimotorically guided* (Orgs et al., 2016; Stevens et al., 2000). Indeed, it relies on SOAs that are sufficiently slow to be consistent with the actual or simulated movement duration (Shiffrar & Freyd, 1990), and thus, sufficient timing might allow for bodily constraints and intuitive physics to influence LAMP. However, two main theories to account for the nature of the influences on LAMP are currently debated in the literature: the *motor* and the *visual theory of LAMP*.

According to the *motor theory of LAMP*, the perception of physically possible movements is grounded in the observer’s sensorimotor representations acquired through motor experience (Funk et al., 2005; Orgs et al., 2016; Saetta et al., 2018; Stevens et al., 2000; Thornton, 1998). It is assumed that fronto-parietal networks tuned to motor control are also involved in the understanding of others’ observed actions (“motor resonance”, Fadiga et al., 1995), which can also be assumed for the LAMP phenomenon. In line with this assumption, the perception of physically possible movements induced by slow SOAs triggers the selective activation of motor areas representing the repertoire of the observer’s possible movements, such as the premotor and primary motor cortices (Orgs et al., 2016; Stevens et al., 2000). Behavioral evidence supporting the motor theory comes, for instance, from studying phantom limb awareness. Experienced by almost all individuals with an amputation (Ramachandran & Hirstein, 1998), phantom limb awareness refers to the persistence of the motor and postural representations of a limb despite its physical absence (Brugger, 2006). The phantom limb might be intentially moved to various degrees or completely immobilized (Saetta et al., 2020a). Training to execute impossible movements with a phantom limb has shown to enhance the visually guided perception of physically impossible movements, regardless of the SOAs (Moseley & Brugger, 2009). Furthermore, individuals with an amputation who typically experience a phantom limb to fade away or bend back once it crosses hindering solid objects (“obstacle shunning”) are more likely to perceive a limb to move straight through an object in the LAMP task (Saetta et al., 2018). Taken together, these findings highlight a tight interrelation between the motor capabilities of a (phantom) limb and the LAMP phenomenon.

Conversely, the *visual theory of LAMP* states that the perception of physically possible movements merely relies on visual perception, and exclusively engages the observer’s visual system (Vannuscorps & Caramazza, 2016). In this view, the extra-striate visual cortex processes static information on human bodies (fusiform body area) or specific body parts (extra-striate body area) (Downing et al., 2001; Peelen & Downing, 2007; Vangeneugden et al., 2014), while the posterior aerea of the superior temporal sulcus reconstructs the movement kinematics (Blake & Shiffrar, 2007b; Grosbras et al., 2012; Puce & Perrett, 2003). In support of this hypothesis, no movement trajectory differences on the LAMP were found between able-bodied controls and individuals with congenital upper limb dysmelia who reported no phantom limb awareness (Vannuscorps & Caramazza, 2016). This suggests that almost complete deprivation of limb motor representations has no impact on LAMP. However, other evidence deriving from the LAMP task revealed that a complete congenital amelia (without accompanying phantom percept) is accompanied by a bias towards a consistently more visually guided perception of physically impossible movements for all SOAs (Funk et al., 2005).

Here, we tested these two divergent theories in two independent studies. In the first study, we used transcranial direct current stimulation (tDCS) in healthy participants to causally interfere with cortical motor processing during the LAMP task. Increasing evidence shows that the cathodal stimulation of the primary motor cortex (M1 c-tDCS) induces hyperpolarisation of the resting membrane potential of the motor cortex’s neurons, reducing its spontaneous activity and excitability (Bindman et al., 1964; Nitsche & Paulus, 2000; Nitsche et al., 2003). Furthermore, a sufficiently long stimulation (duration above 10 min) is accompanied by after-effects lasting up to two hours (Jamil et al., 2017). Previous studies using M1 c-tDCS found reduced M1 activation during action observation and action execution (Qi et al., 2019). In support of the motor theory, we expected that dampening of the primary motor cortex’s activity would reduce the motor simulation of the perceived illusory movements and thus bias LAMP towards the visually guided perception of physically impossible movements. This was expected specifically for slow SOAs.

The second study was conducted in an online (web-based) setting and investigated LAMP in clinical samples presenting atypical alterations in sensorimotor limb representation. We investigated a) individuals with an acquired lower limb amputation and b) individuals affected by body integrity dysphoria (BID) with a desire for, but not yet performed, lower limb amputation. BID is a rare and aberrant condition characterized by dissatisfaction with a normal body morphology or functionality in non-psychotic individuals (Brugger et al., 2016; Saetta et al., 2020b). In its most common form, one limb can be experienced as nonbelonging despite regular anatomical development and integrity of sensory and motor functions. This feeling that a limb does not belong to oneself often leads to the desire for its amputation. Only very recently, in the release of the 11th Revision of the International Classification of Diseases (*ICD-11 — Mortality and Morbidity Statistics*), has BID been recognised as an official mental disorder. A distinctive behaviour displayed by the majority of BID individuals (to varying extent) is the so-called *pretending behaviour*, i.e., the mimicking of the desired body state resembling that of an amputee by moving in a wheelchair, using crutches, or binding up the disowned leg to obtain transient relief of symptoms. A sample of naïve, able-bodied individuals was also included.

Crucially, in all participants, we modulated the actual sensorimotor state by assessing LAMP twice in two different bodily states. Figure 1 describes the three included groups and illustrates the manipulation of the bodily state as implemented in study 2. In support of the motor theory, we generally expected the sensorimotor states to influence LAMP exclusively for slow SOAs (see below for more specific hypotheses).

**Figure 1.**
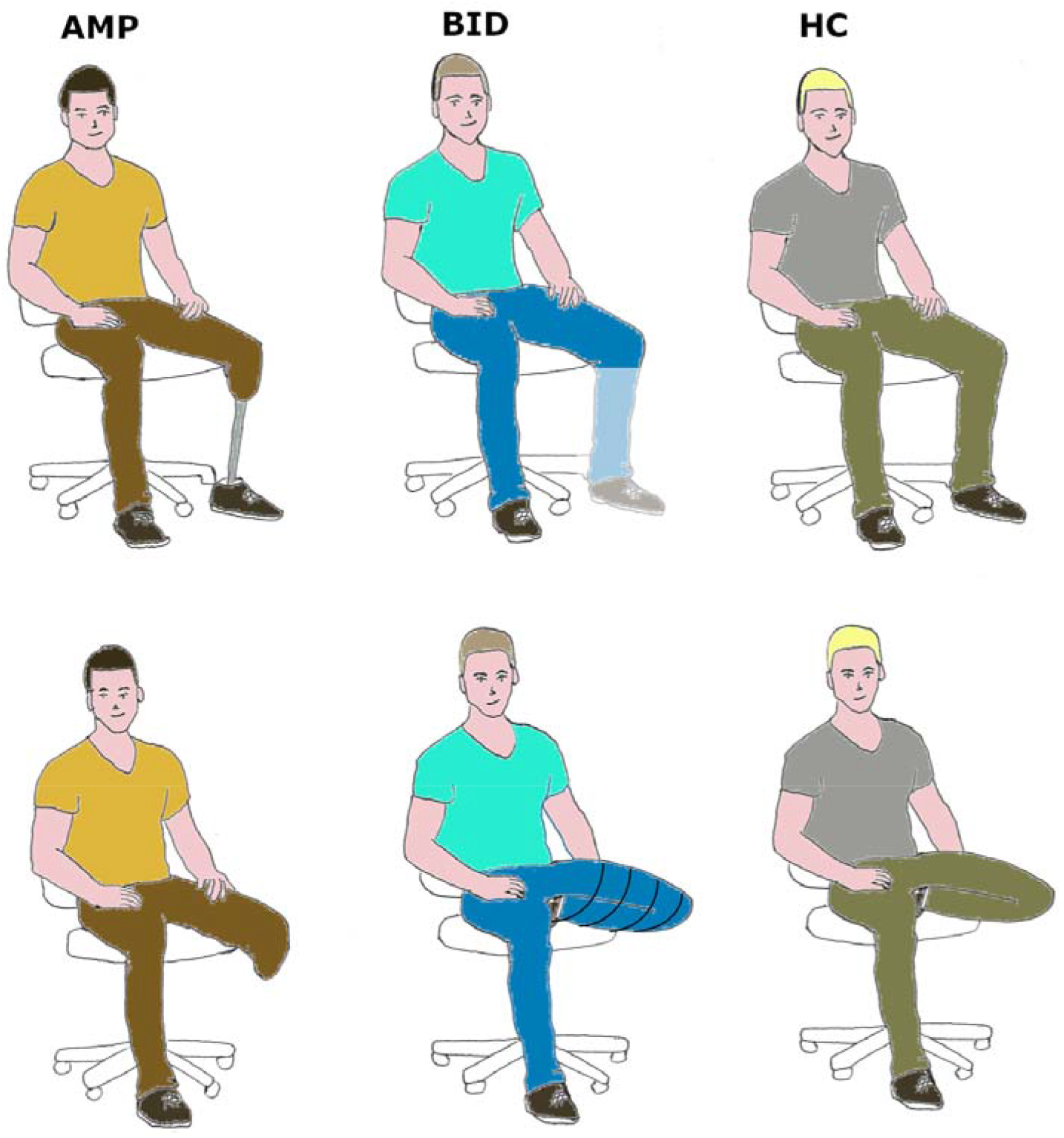
Included groups and sensorimotor state manipulation. Individuals with an amputation (AMP) performed the task twice, either with or without wearing their own prosthesis; Individuals with Body Integrity Dysphoria (BID) sitting in a normal position or pretending (binding up the to-be removed leg; the transparent part of the leg indicates the area for which the amputation desire is reported); Able-bodied individuals (HC) while sitting in a normal position or sitting on one of their legs.

*Individuals with an amputation* performed the task twice, either with or without wearing their own prosthesis (counterbalanced order). Additionally, we measured the integration of the prosthesis into the sensorimotor system, i.e., the subjective sense of prosthesis ownership, defined as the feeling of a prosthetic limb constituting an integral part of the body (Niedernhuber et al., 2018). Accumulating evidence shows that prosthesis ownership positively correlates with prosthesis use (Bekrater-Bodmann et al., 2021). On the other hand, prosthesis use seems to counteract the effects of sensorimotor deprivation from limb amputation, and may drive adaptive plasticity in the sensorimotor cortex (van den Heiligenberg et al., 2018). We expected that higher prosthesis ownership would predict greater bias towards the sensorimotor guided perception of physically possible movements on the LAMP task. This would be observed exclusively for slow SOAs, and only when participants performed the task while wearing the prosthesis.

*BID individuals*, performed the LAMP task twice, while sitting in a normal position or while pretending. We expected that mimicking the desired amputated state would bias LAMP towards the more visually guided perception of physically impossible movements, exclusively for slow SOAs.

*Able-bodied participants* in study 2 completed the experiment twice, either while sitting in a normal position or while sitting on one of their legs. On the basis of previous findings (cp. e.g., showing that restriction of body movements may affect motor control and different cognitive processes (Ionta et al., 2007, 2012)), we expected the peripheral and transient reduction of motor capabilities to bias LAMP towards the more visually guided perception of physically impossible movements, exclusively for slow SOAs.

## Study 1

### Participants

Twenty-four right-handed participants (Males: 13, Females: 11, *M* age = 25.96 years, *SD* = 5.34) with no history of any psychiatric and neurological disorders took part in this study. Exclusion criteria were: presence of epilepsy or seizure, fainting spell or syncope, head trauma, metal implants in the brain/skull, cochlear and neurostimulator implants, cardiac pacemaker, use of recreational drugs, or consumption of more than 3 units of alcohol in the past 24 hours. All participants had normal or corrected-to-normal vision. Participants received monetary compensation for their participation. All participants were informed about the scope of the study and provided written informed consent, complying with the

Declaration of Helsinki (1984), and the approval of the local ethics committee of Utrecht University (protocol number: FETC 19-204)

### Materials

Pairs of photographs, varying only in the position of one of the model’s limbs, were presented. In one photograph, the limb was on the right, and in the other, on the left side of a solid object. In order for participants to select which of the two movement trajectories they perceived, the aforementioned two pictures were superimposed on one another to create a third picture, which showed two arrows depicting the two possible motion trajectories; i.e., the short and physically impossible (i.e., the limb moved through the object) or the long and physically possible (i.e., the limb moved above and around the drawer) trajectory. Fig. 2 represents a sample stimulus pair and the respective superimposed images for the response. The experiment consisted of 96 trials. The pairs of photographs featured left (n = 48) and right (n = 48) upper (n = 48) and lower (n = 48) limbs, and specifically four body parts: i) hand (n = 24), ii) forearm (n = 24), iii) foot (n = 24), iv) shank (n = 24). The model’s limbs were pictured in the first-person (1pp, n = 48) or the third-person (3pp, n = 48) perspective. All these conditions were considered for two reasons: i) to counteract participants’ potential boredeness, making the task more interesting to perform; ii) given a sufficient number of participants, we wanted to explore the effects of previously investigated factors such as limb, laterality, and perspective (Saetta et al., 2018, Saetta et al., in preparation) and their interaction with the newly introduced factor tDCS stimulation.

**Figure 2.**
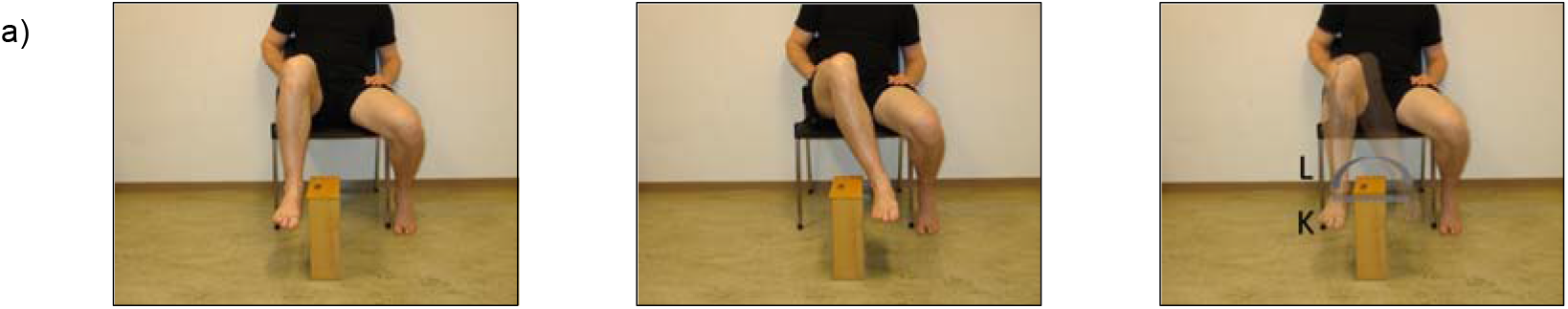
Sample stimulus pair inducing the perception of leg moving through an object, or around the object depending on the SOA (illustrated here is a third person perspective). After the presentation of the stimulus pair, a picture showing two arrows depicting the two possible motion trajectories is presented. The arrow “L” indicates the more sensorimotor guided perception of physically possible movements. The arrow “K” the more visually guided perception of physically impossible movements.

### Procedure

In a single-blind, sham-controlled between-subject design, participants were assigned to either the real stimulation condition, where cathodal tDCS was applied over the left motor cortex (experimental stimulation; n=12, Males: 6, Females: 6, *M* age = 27.58 years, *SD* = 7.09) or the sham stimulation (n=12, Males: 7, Females: 5, *M* age = 24.33 years, *SD* = 1.87), according to a Latin Square counterbalancing assignment to conditions. Given the role of the left hemisphere in initiating bimanual movements (Walsh et al., 2008), and that inhibition of the ipsilateral motor cortex has been shown to affect both the ipsilateral and contralateral hand movements (review in Chen et al., 1997), we expected that stimulation of the left motor cortex would extert bilateral effects, rather than a specific effect for the contralateral limb. There was no statistically significant age difference between groups (two-tailed t(12.53) = 0.154, p = 0.15). The stimulation was delivered with a battery-driven, constant-current stimulator (neuroConn DC-stimulator) through 35 cm^2^ sponge electrodes on which the Ten20 conductive paste was spread. The voltage was set to 1 mA. In the experimental stimulation condition, the tDCS was applied for 20 minutes. The sham stimulation consisted of 20 minutes in total, with only the first 5 sec applying tDCS stimulation, after which the stimulation was deramped for 10 sec. The procedure lasted for 20 minutes irrespective of the stimulation condition (experimental or sham), and all participants perceived the current flow as an itching sensation, as they confirmed verbally.

The electrodes were placed over the C3 and Fp2 according to the 10/20 EEG system (Jasper 1957), with the cathode being placed over the left motor cortex (Nitsche & Paulus, 2001) and the anode over the contralateral orbital/supraorbital region. The COMET toolbox (Jung et al., 2013) implemented in MATLAB was used to simulate the electric field generated by the tDCS with the present electrodes configuration and size. This configuration has proven effective in down-regulating motor cortex excitability in replicated and multi-approach studies combining tDCS and single-pulse or repetitive transcranial magnetic stimulation (TMS) (Nitsche & Paulus, 2000; Siebner et al., 2004). After the stimulation, participants were familiarized with the task via five practice trials before the actual experiment started.

Instructions were provided verbally by the experimenter. Additionally, before the start of the experiment, written instructions appeared at the center of the screen. A special emphasis was placed on explaining that there were no wrong or correct responses, as perception is highly subjective. Participant were asked to respond as quickly as possible and to not simulate the observed movement, but rather to remain relaxed in the sitting position. The distance from the screen was set to approximately 50 cm. The experiment was programmed in E-prime, version 3.

In each trial, after a central fixation cross was shown for 1 sec, the pairs of photographs were presented alternately with different stimulus durations (StimD) and interstimulus intervals (ISI) at five SOAs. As in Shiffrar & Freyd (Shiffrar & Freyd, 1990), the shortest SOA was 150 msec (SD = 100 msec, ISI = 50 msec). The other SOAs were 250 msec (SD = 200 msec, ISI = 150 msec), 450 msec (SD = 400 msec, ISI = 350 msec), 650 (SD = 600 msec, ISI = 550 msec), and 750 msec (SD = 700 msec, ISI = 650 msec). The experiment consisted of two blocks: one where the actor’s movements were observed from 1pp and the other were they were observed from 3pp. The factor perspective was introduced: On the basis of previous studies (Ruby & Decety, 2001), we expected stimuli in 1pp, compared to 3pp, to more likely elicit a motor simulation of the observed movements, and therefore to bias LAMP towards the more sensorimotorically guided perception than stimuli in 3pp, which are associated with encoding processes in the observer’s visual system and would therefore bias LAMP towards the more visually guided perception. The presentation of the blocks was counterbalanced across the participants. The experiment lasted approximately 50 minutes in total.

### Data Analysis

Data were analysed with R Studio v. 1.3.1093. The outcome measure, *the illusion experience*, was defined as the tendency to report physically impossible movements (coded as −1) or physically possible movements (coded as +1).). The R scripts and the dataset are deposited on the open science framework (OSF, https://osf.io/4z6yc/). Linear mixed models considering the most relevant factors according to our hypotheses were fitted with the R lme4 package (Bates et al., 2015 They were estimated using REML and nloptwrap optimizer. 95% Confidence Intervals (CIs) and p-values were computed using the Wald approximation. The appropriateness and the need for a linear mixed model were assessed by applying the methods reported in Field et al. (2012). A formal comparison of two models, one including the intercept, and the other allowing the intercept to vary across the participants, was performed by examining the changes in the −2log-likelihood (Bliese & Ployhar, 2002). The model’s fit significantly improved as a result of setting a random intercept for each participants (X^2^(1) = 212.95, p < 0.001). A linear mixed model examined the illusion experience as a function of the factors tDCS stimulation (experimental/sham) and SOA (150, 250, 450, 650, 750) and the participant’s responses. The model was the following:

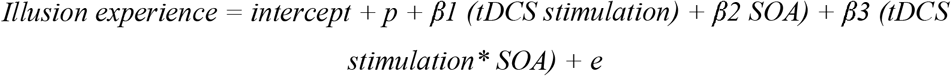

where “*β_x_*” stands for the estimated parameters, “*e”* stands for the residuals, and “*p*” represents the random intercept for participants. A type III ANOVA with Satterthwaite’s method table was produced to detect the main and the interaction effects. To explore the significant effects, post hoc pairwise analyses were performed with the emmeans package (Lenth, 2020). Degrees of freedom (df) were calculated with the Kenward-Roger method. Bonferroni correction for multiple comparisons was applied. Further analyses exploring the effects of laterality are reported and discussed in the supplementary materials.

### Results

With this procedure, a notable number of observations (n = 2304) was modelled while adjusting for the within-subject and within-group dependency, resulting in substantial explanatory power (conditional R^2^ = 0.34) for the model. Results show a significant main effect of tDCS stimulation (F(1,22) = 5.80, p = 0.02), indicating that participants were more likely to perceive physically impossible movements in the experimental M1 c-tDCS stimulation, as compared to the sham stimulation condition, and a significant main effect of SOA (F(4,2272.5) = 176.30, p < 0.0001), indicating that with slower SOAs, participants were more likely to perceive physically possible movements. Means and standard deviations for the combination of the levels of the factors tDCS stimulation and SOAs are reported in table 1 in the supplementary materials. Post hoc tests showed significant differences for the comparison between SOA_150_ and SOA_250_ (*Mdiff =* −0.32*, SE =* 0.05 *, df =* 2273*, t.ratio =* - 5.84, p < 0.0001), and SOA_250_ and SOA_450_ (*Mdiff =* 0.65, *SE =* 0.05, *df* = 2272, *t.ratio =* - 12.022, p < 0.0001), but not for the comparison between SOA_450_ and SOA_650_ (p = 0.16), and SOA_650_ and SOA_750_ (p = 1). Furthermore, a significant interaction of tDCS by SOA (F(4, 2272.5) = 4.68, p < 0.0001) was observed. Post hoc tests showed no significant differences in the illusion experience between the experimental and the sham tDCS (contrast: experimental tDCS – sham tDCS) for the fastest SOA_150_ (*Mdiff =* −0.05 *, SE =* 0.08 *, df =* 34.81*, t.ratio =* - 0.64 *, p =* 0.53) and SOA_250_ (*Mdiff =-*0.11 *, SE =* 0.08 *, df =* 34.77*, t.ratio =* −1.41 *, p =* 0.17). However, significant differences were found for the slower SOAs; SOA_450_ (*Mdiff =* - 0.21*, SE =* 0.08 *, df =* 34.80*, t.ratio =* −2.77 *, p =* 0.009), SOA_650_ (*Mdiff =* −0.19*, SE =* 0.08 *, df =* 34.68*, t.ratio =* −2.55 *, p =* 0.015), SOA_750_ (*Mdiff =* −0.25 *, SE =* 0.08 *, df =* 35.40*, t.ratio =* −3.35 *, p =* 0.002). That is, participants were more likely to perceive physically impossible movements after experimental M1 c-tDCS than after sham stimulation, but this stimulation-dependent effect was only observed when stimuli were presented at the slower SOAs. Results are visualized in Figure 3.

**Table 1.** Questions asked to individuals with lower limb amputation with the type of scales and the scale values.

| Item | Question | Scale type | Scale Values |
| --- | --- | --- | --- |
| Frequency of prosthesis use | How often do you use your prosthesis in a week? | ordinal | 0 = not at all, 1 = less than twice, 2 = every second day, 3 = almost daily, 4 = daily |
| Prosthesis ownership | How much do you perceive your prosthesis as part of your body when you wear it? | continuous | 0 = prosthesis feels like a foreign body – 10 = prosthesis is like a part of the body |
| Prosthesis sense of control | How much control do you have over the movements of your prosthesis when you wear it? | continuous | 0 = I have no control – 10 = I have full control) |
| Phantom limb awareness (last 3 months) | Have you recently (over the past three months) experienced the presence of a phantom limb? | categorical | 0 = No, I have never experienced a phantom limb, 1 = No, but I used to experience a phantom in the past, 2 = Yes, I currently am experiencing a phantom limb |
| Phantom limb awareness strength (last 4 weeks) | How strong were these experiences on average over the past four weeks? | continuous | 0 = not present – 10 = very strong |
| Phantom limb awareness frequency | How often do you feel the phantom limb? | ordinal | 0 = never, 1 = less than once a month, 2 = once a month, 3 = every other week, 4 = 1-2 a week, 5 = at least 3 days a week, 6 = at least 5 days a week, 7 = once a day, 8 = several times a day, 9 = all the time |

**Figure 3.**
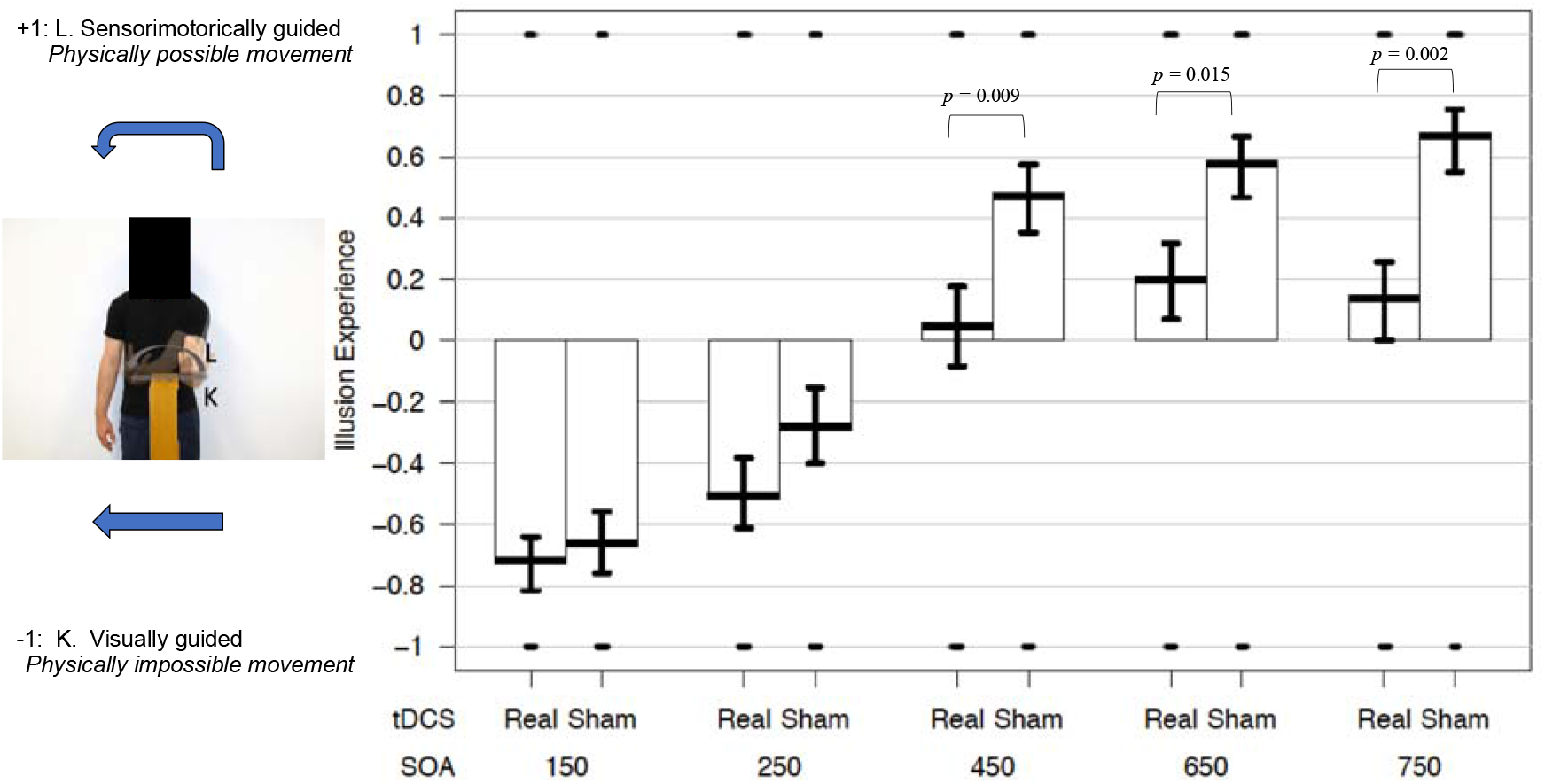
Illusion experience in the LAMP task. Participants who underwent the experimenteal M1 c-tDCS compared to those who underwent the sham stimulation were biased toward the visually guided perception of physically impossible movements after the stimulation. This stimulation-dependent effect was only observed when stimuli were presented at the slower SOAs. Mean values for each condition and standard errors are reported.

### Discussion study I

In a sham-controlled M1 c-tDCS study, we confirmed the contribution of sensorimotor representations to the LAMP by interfering with the cortical activity and excitability of the primary sensorimotor areas, thereby providing support for the motor theory. M1 c-tDCS has previously been shown to induce long-term depression effects in the sensorimotor cortex (Nitsche & Paulus, 2000). In a recent study, a reduced amplitude of motor evoked potentials due to M1 c-tDCS was registered in connection with action observation and excecution (Qi et al., 2019). Another study found that M1 c-tDCS lowers the accuracy of predictions of partially occluded human, but not non-human, reaching-grasping movements (Paracampo et al., 2018). These studies showed that motor theory applies to action observation, execution, and prediction, as revealed by the inhibitory effects of M1 tDCS on all these processes. Our results extend these findings by showing an effect of M1 c-tDCS on the LAMP phenomenon. As in previous studies (Saetta et al., 2018; Shiffrar & Freyd, 1990; Stevens et al., 2000), we found that fast SOAs biased LAMP towards the visually guided perception of physically impossible movements. Slow SOAs instead induced the perception of physically possible movements, which are, based on previous findings (Orgs et al., 2016; Stevens et al., 2000), sensorimotorically guided. More importantly, M1-tDCS biased LAMP towards more visually guided perception. This effect was specific for the slow SOAs. While previous studies applied correlational methods, such as positron emission tomography (Stevens et al., 2000, PET) and functional magnetic resonance imaging (Orgs et al., 2016, fMRI), we here implemented a causative method to corroborate the motor theory of the LAMP for the first time (to the best of our knowledge).

## Study 2

### Participants

Twenty-nine individuals with unilateral lower limb amputation participated in the online study (Men: 24, Women: 5, Right-sided amputation: 12, Left-sided amputation: 17, Mean age = 52.66 years, SD = 5.09). These individuals were recruited using the PHANTOMMIND data base (first description by (Bekrater-Bodmann et al., 2015)). Inclusion criteria for the present study were a) acquired major lower limb amputation, b) age between 18 and 80 years, and c) using a prosthesis. Frequency of prosthesis use, prosthesis ownership, and sense of control over the prosthesis, as well as presence, frequency, and strength of phantom limb awareness over the last four weeks were assessed using a questionnaire. Table 1 reports used questions, the scale types, and the scale values.

Individuals with an amputation used their prosthesis with a high frequency (Median frequency of prosthesis use = 4 per week (scale ranging from 0 to 4, MAD = 0, see table 1), indicating daily use. Overall, prosthesis ownership (Mean = 7.45, SD = 2.63) and sense of control over the prosthesis (Mean = 8.38, SD = 1.40) were rated as high. Nineteen individuals with an amputation reported current phantom limb awareness, 5 had experienced a phantom limb in the past, and 5 had never experienced a phantom limb.

Ten individuals with BID were recruited through online forums or personal contacts established in previous studies (Saetta et al., 2020b) (Males: 8, Females: 2, Right-sided amputation desire: 7, Left-sided amputation desire: 3, Mean age = 41.00 years, SD = 12.36). The desire for amputation was assessed using the Zurich Xenomelia Scale (Aoyama et al., 2012). This 12-item scale consists of 3 subscales assessing different aspects: “amputation desire” (identity restoration as the main motivation for the amputation), “erotic attraction” (sexual arousal to amputated bodies), and “pretending” (inclination to mimic an individual with amputation). The subscores for each scale are the sum of 4 item scores. The subscore can range from 1 = not intense to 24 = most intense. The average subscores were: amputation desire: Mean = 22.70, SD = 1.70, erotic attraction: Mean = 16.70, SD = 6.07, pretending: Mean = 17.00, SD = 2.49.

Thirteen naïve able-bodied men (Mean age = 40.54 years, SD = 6.36), not included in study 1, were recruited through the platform “Prolific” (https://www.prolific.co/).

All participants gave their informed consent. The informed consent was displayed prior to starting the online survey, and participants had to actively give there consent by ticking a box. The study was approved by the Local Ethics Committee of the Faculty of Arts and Social Sciences at the University of Zurich (Approval number: 17.12.8).

### Materials

Participants were confronted with pairs of photographs varying only in the position of one of the model’s limbs. In one condition, as in study 1, the limb was once on the right and once of the left side of an object (object solidity condition). Additionally, in another condition, in one photograph the limb was on the right and in the other on the left side of the contralateral limb (limb solidity condition; see Fig 2b). As in study 1, for response selection, the two pictures of a pair were superimposed onto one another to create a third picture that showed two arrows depicting the two possible motion trajectories (i.e., limb moving through or around the object). This was done so participants could select their perceived movement trajectory.

The experiment consisted of the presentation of 256 stimulus pairs. They featured left (n = 128) and right (n = 128), upper (n = 128) and lower (n = 128) limbs, and specifically four body parts: i) hand (n = 64); ii) forearm (n = 64), iii) foot (n = 64), iv) shank (n = 64). The model’s limbs were pictured in the first-person (1pp, n = 128) or third-person (3pp, n = 128) perspective, and belonged to the object solidity constraint (n = 128) or limb solidity constraint (n = 128) condition. These two conditions were included on the basis of two strains of evidence: i) whether individuals with amputations show obstacle shunning or not depends on whether the phenomenal space occupied by the phantom limb overlaps with that of biological (i.e., the contraleral limb) or non biological (a wall) matter (Saetta et al., 2020a), and ii) obstacle shunning experience is related to LAMP (Saetta et al,. 2018).

### Procedure

The jsPsych software (https://www.jspsych.org) (de Leeuw, 2015) was used to program the online experiment. A trial consisted of a central fixation cross shown for 1 sec, followed by 256 stimulus pairs that were presented alternately with different stimulus duration (StimD) and interstimulus interval (ISI) at four SOAs. The shortest SOA was 150 msec (SD = 100 msec, ISI = 50 msec). The other SOAs were 250 msec (SD = 200 msec, ISI = 150 msec), 650 (SD = 600 msec, ISI = 550 msec), and 750 msec (SD = 700 msec, ISI = 650 msec). The experiment consisted of a single block, in which all the trial types were randomised. The duration of the experiment was approximately 15 minutes.

Before the start of the experiment, as in study 1, all the instructions were presented at the center of the screen. Participants’ sensorimotor states were manipulated by performing the experiment twice (in individuals with an amputation: either while wearing a prosthesis or while not wearing a prosthesis; in BID individuals: either while binding up their unwanted leg (pretending) or while sitting in a normal position; in able-bodied participants: either while sitting on one of their legs or while sitting in a normal position, see Figure 1). Four participants sat on the right leg, 9 on left leg. For each group, the order of conditions was counterbalanced. The two assessments were performed on two separate days, with a mean delay of 7 days.

### Data analysis and results

Linear mixed models were fitted following the statistical methods described for study 1. R Studio v. 1.3.1093 was used. The R scripts and the dataset are deposited on the open science framework (OSF, https://osf.io/4z6yc/). Separate analyses were conducted on the three samples.

#### Individuals with an amputation

A linear mixed model examined the impact of the factors sensorimotor state, SOA, and prosthesis ownership (a continuous variable) on the illusion experience. This model also included the two-way and the three-way interactions between these parameters. A random intercept for each participant was set given the random structure of the data (X2(3) = 659.25, p < 0.001). The number of the modelled observation was 7424 and the model’s total explanatory power was moderate (conditional R^2^ = 0.19). Results are reported in table 2.

**Table 2.** Results of the linear mixed model examining the impact on the illusion experience in the LAMP task in individuals with a lower limb amputation of sensorimotor state, SOA, and prosteshis ownership.

|  | Sum Sq | Mean Sq | NumDF | DenDF | F value | Pr(>F) |
| --- | --- | --- | --- | --- | --- | --- |
| Sensorimotor state | 0.03 | 0.03 | 1 | 25 | 0.04 | 0.842 |
| SOA | 36.86 | 12.29 | 3 | 7383 | 14.58 | <0.0001 |
| Prosthesis ownership | 0.41 | 0.41 | 1 | 25 | 0.49 | 0.49 |
| Sensorimotor state*SOA | 6.42 | 2.14 | 3 | 7383 | 2.54 | 0.055 |
| Sensorimotor state*prosthesis ownership | 0.30 | 0.30 | 1 | 25 | 0.36 | 0.554 |
| SOA*prosthesis ownership | 6.05 | 2.02 | 3 | 7383 | 2.39 | 0.067 |
| Sensorimotor state*SOA*prosthesis ownership | 13.77 | 4.59 | 3 | 7383 | 5.45 | <0.0001 |

The results show a main effect of SOA (F(1,7383 = 14.58, p < 0.0001). Post hoc tests showed significant differences for the comparison between SOA_150_ and SOA_250_ (*Mdiff =* - 0.09*, SE =* 0.03 *, df =* 7383*, z.ratio =* −2.83, p = 0.028), and between SOA_250_ and SOA_650 (_*Mdiff =* −0.24, *SE =* 0.03, *df* = 7383, *z.ratio =* −7.53, *p* <.0001), but not for the comparison SOA_650_ and SOA_750_ (p = 1). We also found a three-way interaction effect of sensorimotor state by SOA by prosthesis ownership (F(3,7383 = 5.45, p = 0.001). As shown in Figure 4, the more the prosthesis was felt as part of the body, the more individuals with an amputation were likely to perceive a physically possible movement, but only while *wearing the prosthesis*. This effect was observed exclusively for the slowest SOAs, as shown by the continuous and the stippled lines representing the two sensorimotor states intersecting, and having slopes that are opposite to each other at the SOA_650_ (turquoise lines) and at SOA_750_ (violet lines), but not at SOA_250_ (green lines), and at the SOA_150_ (red lines). Further explorative analyses exploring the effects of limb, consistency of the side of the amputation with the laterality of the stimuli, solidity and perspective are reported in the supplementary materials.

**Figure 4.**
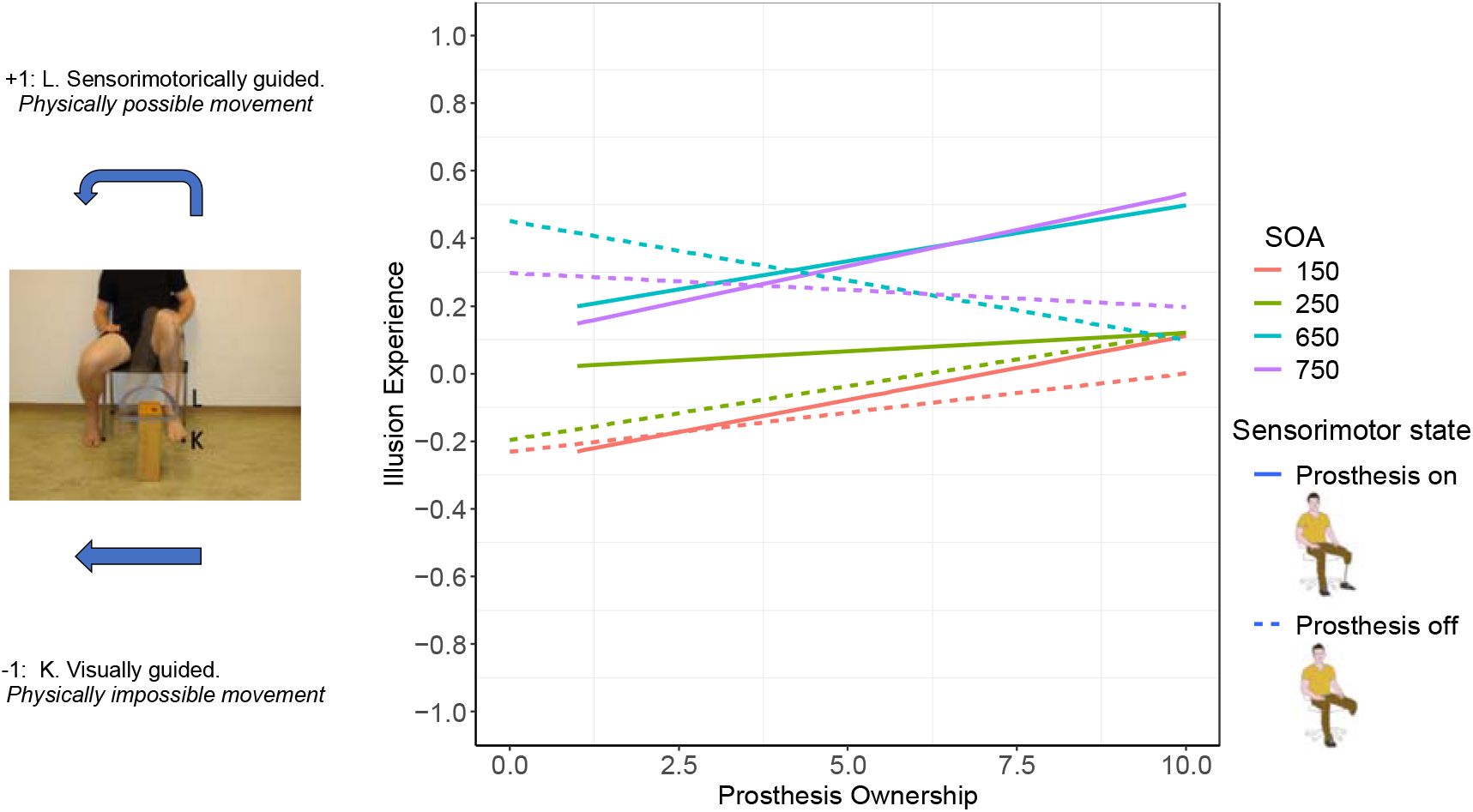
Illusion experience in the LAMP task in individuals with a lower limb amputation. The more the prosthesis was felt as part of the body, the more participants were biased towards the more sensorimotorically guided perception of physically possible movement, but exclusively while they wore the prosthesis compared to when they did not. As predicted, this was only observed when stimuli were presented at the slower SOAs.

#### BID individuals

A linear mixed model examined the factors sensorimotor state and and their interaction in the LAMP task. A random intercept for each participant was set, given the random structure of the data (X2(3) = 404.88, p < 0.0001). The number of the modelled observation was 2560 and the model’s total explanatory power was weak (conditional R^2^ = 0.12). The results show a significant main effect of SOA (F(3, 3308) = 19.40, p < 0.0001). Post-hoc tests showed no significant differences for the comparison between SOA_150_ and SOA_250_ (*p* = 1), significant differences for the comparison between SOA_250_ and SOA_650 (_*Mdiff =* −0.26, *SE =* 0.05, *df* = 2543, *t.ratio =* −5.170, *p* <.0001), but not for the comparison SOA_650_ and SOA_750_ (p = 1). We found an effect of sensorimotor state (F(1, 2543) = 4.47, p = 0.035). The latter effect shows that BID individuals were more likely to perceive physically impossible movements while binding up their leg, than while sitting in a normal postion (see Figure 5). The interaction sensorimotor state by SOA was not significant (F(3, 2543) = 0.54, p = 0.65). Further explorative analyses exploring the effects of limb, consistency of the side of the to-be removed leg with the laterality of the stimuli, solidity and perspective are reported in the supplementary materials.

**Figure 5.**
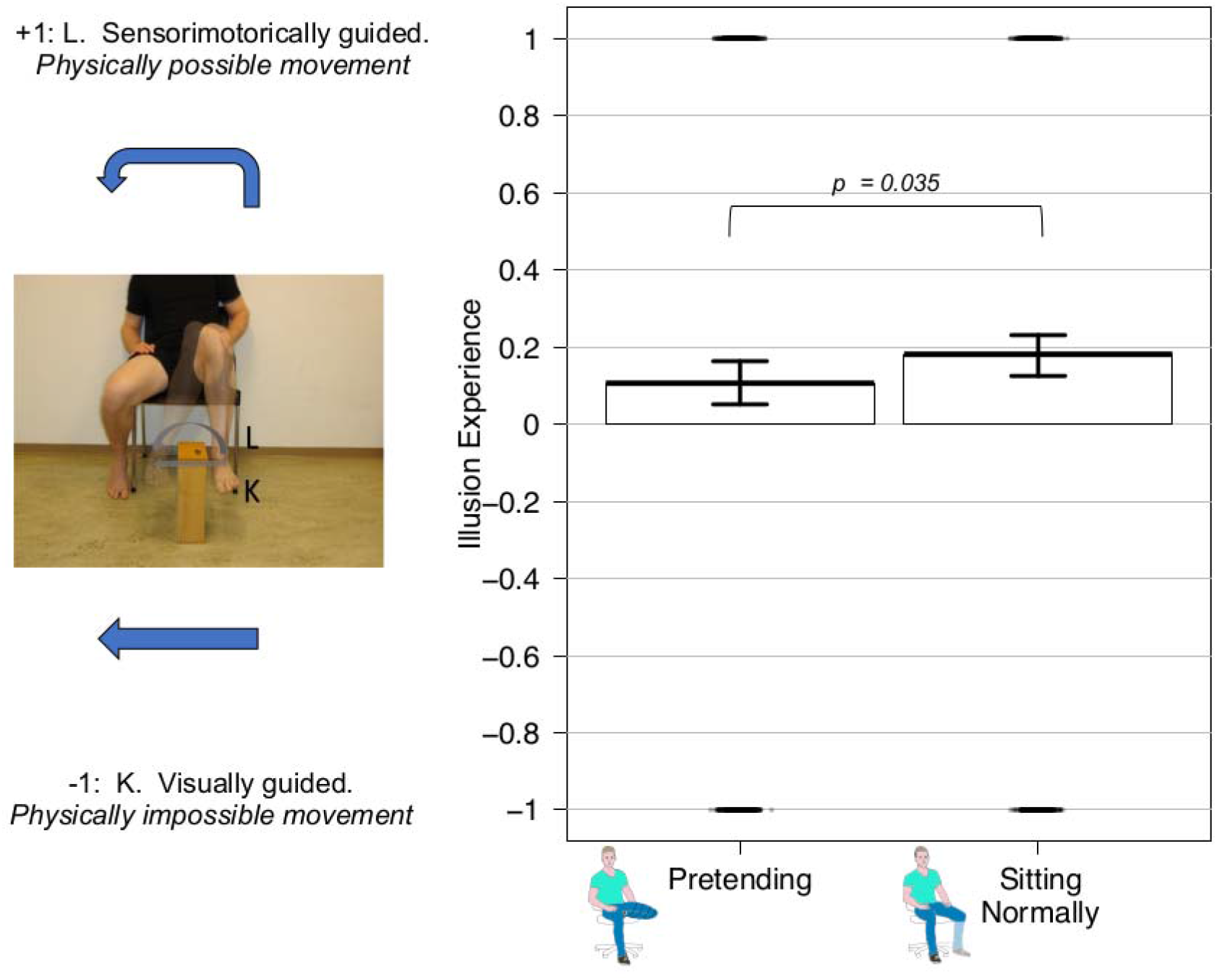
Illusion experience in the LAMP task in BID individuals. Participants were biased towards the visually guided perception of physically impossible movements while binding up their leg compared to while sitting in a normal position. Displayed are means and SEs.

#### Able-bodied individuals

A linear mixed model examined the impact on the illusion experience of the factors sensorimotor state and SOA, and their interaction. A random intercept for each participant was set given the random structure of the data (X2(1) = 135.12, p < 0.0001). The number of the modelled observation was 3328 and the model’s total explanatory power was weak (conditional R^2^ = 0.12). We found a significant main effect of SOA (F(3, 3308) = 71.30). Post-hoc tests showed no significant differences for the comparison between SOA_150_ and SOA_250_ (p = 1), but significant differences for the comparison between SOA_250_ and SOA_650 (_*Mdiff =* −0.49, *SE =* 0.046, *df* = 3308, *t.ratio =* - 10.50, *p* <.0001), but not for the comparison SOA_650_ and SOA_750_ (p = 1). We found no main effect of sensorimotor state (F(1, 3308) = 1.96, p = 0.16) was found. However, the interaction of sensorimotor state by SOA was significant (F(3, 3308) = 5.98, p = 0.0005). Post-hoc tests showed a significant differences in the illusion experience between the the two sensorimotor states (contrast: normal sitting – binding up on the leg) for the SOA_250_ (*Mdiff =* −0.04 *, SE =* 0.06 *, df =* 3308*, t.ratio =* −0.66*, p =* 0.50), and a significant difference for the SOA_150_ (*Mdiff =*-0.12*, SE = 0.06, df =* 3308*, t.ratio =* −1.99, p *=* 0.047). However, significant differences were found for the slower SOAs: SOA_650_ (*Mdiff =* 0.22*, SE =* 0.07 *, df = 3308, t.ratio =.31,* p *= 0.0009*), SOA_750_ (*Mdiff =* 0.14 *, SE =* 0.07 *, df = 3308, t.ratio =* - 2.13*, p =* 0.03). Able-bodied individuals were thus more likely to perceive physically impossible movements when sitting on their leg than when sitting in a normal postion, but only for the slowest SOAs. Results are displayed in Figure 6.

**Figure 6.**
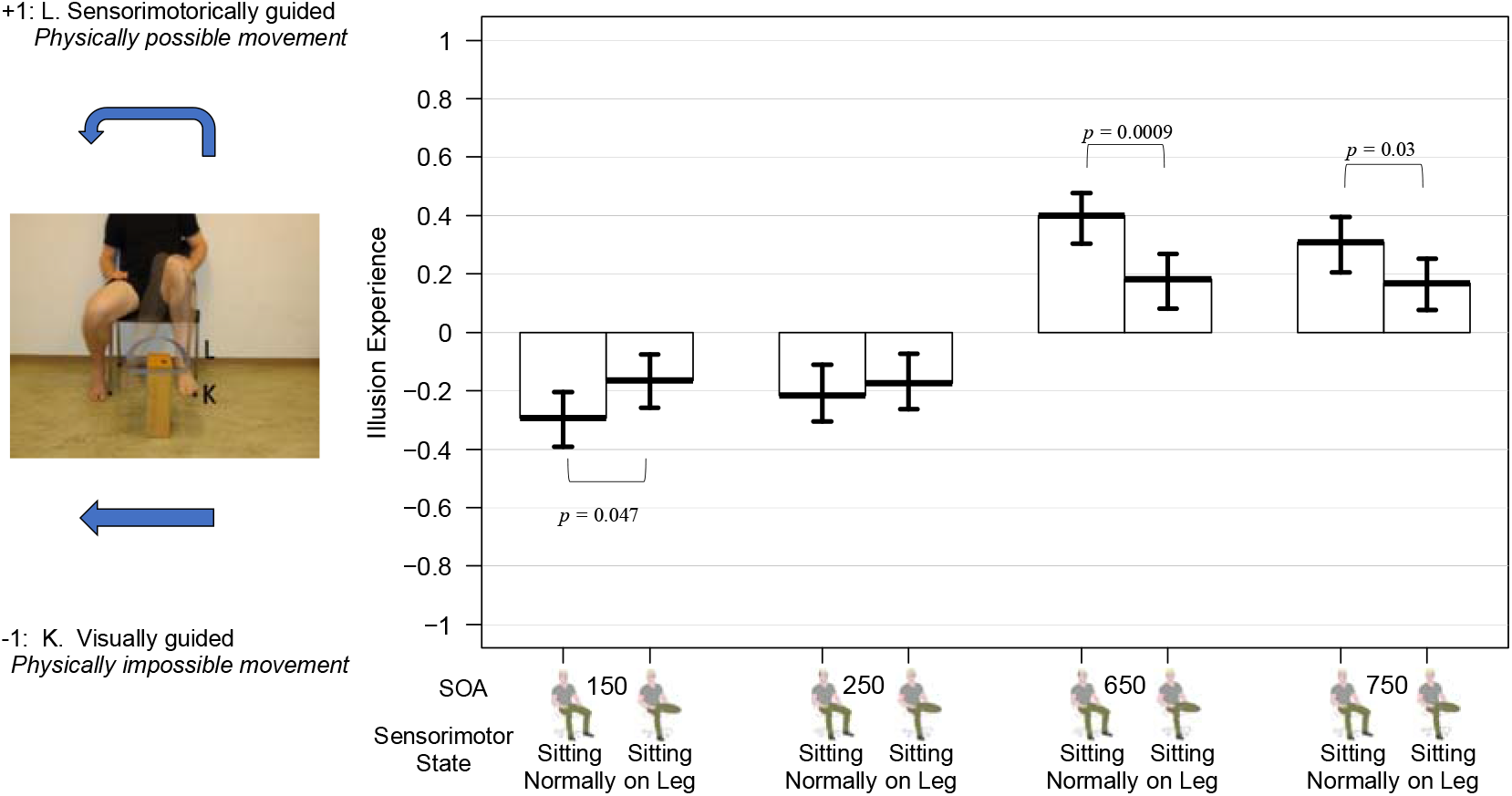
Illusion experience in the LAMP task in able-bodied individuals. Participants were biased towards the visually guided perception of physically impossible movements while sitting on their leg compared to while sitting in a normal position. This was only observed when stimuli were presented at the slower SOAs. Mean values for each condition and standard errors are reported.

Further explorative analyses exploring the effects of limb, consistency of the side of the leg on which participants sat with the laterality of the stimuli, solidity, and perspective are reported in the supplementary materials.

### Discussion Study II

This web-based study revealed an influence of the current (peripherally modulated) sensorimotor states on LAMP in three different participant samples. Specifically, we showed that i) the higher the prosthesis ownership, the more likely individuals with an amputation were to report a sensorimotorically guided perception, when they *wore the prosthesis*, and when the SOAs were slow; b) BID individuals were more likely to report the visually guided perception when pretending (i.e., when mimicking their desired, amputated state by binding up their legs); and c) healthy controls were more likely to show a visually guided LAMP when sitting on their legs, but only for the slow SOAs. While the results for each sample will be discussed in more detail below, our findings generally seem to be in line with the motor theory of LAMP. Additionally, we replicated the well-known effect of the SOA in the LAMP task for the first time in a web-based setting.

Our first finding indicates a modulation of wearing a prosthesis in the perception of the LAMP task that was dependent on the degree of prosthesis ownership. Notably, most individulas with a lower limb amputation wore their respective prostheses daily, and experienced high ownership of their prosthesis. Accumulating theoretical and empirical evidence suggest that the integration of a prosthesis into an amputee’s body representation enhances prothesis use (Bekrater-Bodmann, 2021). Prosthesis use has been identified as a key factor in the development of alternative motor representations adapting to limb loss (van den Heiligenberg et al., 2018). In particular, it has been shown that prosthesis use may recruit the same neural networks that would have normally been recruited by the the limb prior to amputation in large scale visual and motor networks. Notably, this “replacement” of the neural representation by the prosthesis in sensorimotor areas that had hosted the representation of the limb prior to amputation is not dependent on the visual exposure to, or familiarity with, the prosthesis, but specific to its use (van den Heiligenberg et al., 2018). In line with the motor theory, we found that wearing the prothesis, compared to not wearing it, induced more sensorimotor guided perceptions in the LAMP task. Notably, this bias was dependent on the degree of prosthesis ownership and was specific for slow SOAs. Our results suggest that for the LAMP task, a prosthesis that is felt as part of the body and is frequently used poses the same physical constraints as the intact limb does (see also below, but see Bekrater-Bodman et al., 2021, who caution the simplifying equation of “embodiment” and “frequency of use” of a prosthesis). Future studies combining fMRI with the LAMP task should look at the relationship between LAMP and the neural activation related to prothesis perceptions to verify this link on a neural level.

The second finding showed an effect of the bodily posture on perception in the LAMP task in BID individuals. Pretending (i.e., simulating the desired amputation) constituted one of the bodily states. While petending allows for the temporary alignment of the desired and actual body, and provides an instant and transient relief from the desire for amputation (First, 2005), its repetition over time may effect a shrinking of the corresponding sensorimotor cortical representations due to immobilization. Indeed, Liepert et al. (1995) showed, in able-bodied participants, a linear relationship between the duration of lower limb immobilitation and shrinking of the spatial extent of the leg sensorimotor representation, as measured by TMS. Relatedly, long-term immobilization has been shown to induce significant structural brain changes, such as a reduction of cortical thickness in the primary sensorimotor cortex of the corresponding limb (Langer et al., 2012). Futhermore, limb disuse has been shown to be accompanied by reduced neural activity in the corresponding primary motor cortex (Gandola et al., 2017). In accordance with these findings, we recently showed a negative correlation between pretending and the concentration of gray matter in the right superior parietal lobule (Saetta et al., 2020b), a key node for both motor execution and motor imagery networks (Solodkin et al., 2004). Along these lines, we showed that BID individuals in their pretending state were biased towards the more visually guided perception on the LAMP task. While we expected this effect based on previous literature to be specific for slow SOAs, this was evident only when collapsing across all the fast and slow SOAs.

The third finding was that able-bodied controls were also biased towards the more visually guided perception during transient immobilitazion of the leg (i.e., while sitting on it). This pattern was presumably observed due to a reduction of peripheral sensorimotor signalling (cp. e.g., Ionta et al., 2007, 2012). Contrary to our findings in BID individuals, and in line with our hypothesis, the effect in the able-bodied individuals was specific for the slow SOAs. The results are consistent with the motor theory, and consistent with previous findings demonstrating an influence of body posture and peripheral body movement restraint on other motor-related cognitive processes, such as mental rotation of body parts (Ionta et al., 2007) and action perception (Zimmermann et al., 2013).

### General Discussion

Across two studies using a similar LAMP task, and in line with the motor theory, we showed that both central (study 1) and peripheral (study 2) manipulations of sensorimotor states affected LAMP. In study 1, the dampening of the activity of the motor cortex induced a bias towards the more visually guided perception of impossible movements. In study 2, assuming different (clinically relevant) bodily states critically modulated LAMP in a way that was consistent with the observers’ sensorimotor representations accumulated through motor experience. In particular, in individuals with an amputation, the integration of the prosthesis into the sensorimotor system induced the more sensorimotorically guided perception of physically possible movements. In parallel, BID and able-bodied individuals were also biased toward the more sensorimotorically guided perception while performing the task in an “full body” sensorimotor state (in the physical sense). However, when this full body sensorimotor state was affected by not wearing the prosthesis by the transient pheripheral immobilization of the leg by sitting on it, visually guided perception was more likely to occur. We found however a differential influence of SOAs on LAMP while sitting on the leg in BID individuals and able-bodied. We speculate that this might depend on the differences in the accumulated motor experience between these groups that could be due to the pretending in BID individuals.

Both studies come with important limitiations. First, while our sample size in study 1 was comparable to or larger than that of previous studies (Saetta et al., 2018; Shiffrar & Freyd, 1990; Stevens et al., 2000), and the statistical power was adequate, the sample sizes remain modest. Therefore, replication studies in bigger sample sizes would be desirable. Second, M1 tDCS has shown to exert minor or no effects on corticospinal activity in almost 50% of the cases (Wiethoff et al., 2014). As the present study lacks an objective measure of cortical excitability, we cannot accurately infer whether this might have been the case in our sample. However, such a lack of neural modulation should be reflected in a *smaller* effect of the stimulation, and there is no reason to assume confounding effects in our included groups. Third, the effects of M1 tDCS may extend to the neighbouring sensory regions and act on large-scale cortico-subcortical networks (Stagg et al., 2018), making it impossible to accurately disentangle the effects of sensory and motor processes on LAMP. The weaknesses inherent to the M1 tDCS would require the integration of a single-pulse TMS protocol as a more robust technique to target specific regions of M1 and systematically quantifiy the effects of this stimulation on cortical excitability. The use of this technique in future studies may provide useful insight to further substantiate the motor theory.

In study 2, due to its web-based nature, there were no means to control whether the manipulation of the sensorimotor states was in fact realized, other than a manipulation check question asking the participants to answer truthfully whether they had followed the required procedure. Future replications in a laboratory setting would thus be advantageous. However, the non-compliance to the manipulation should have reduced the evinced effects of sensorimotor state rather than enhance them. Furthemore, the replication of the main effect of SOA in all three samples alludes to the validity of the results, and justifies an online application of the task. By administering the task online, we were able to recruit a sample of 29 individuals with an amputation to perform the task twice, resulting in a larger sample size than in classical LAMP studies. Although the number of BID inviduals is admittedly small, the presumed rarity and secrecy of the disorder (First, 2005) often sets hurdles in recruiting an adequately powered sample size. Furthermore, while we were able to follow up with amputees with a reminder in the case of non-participation in the second session a after one week, the anonymous nature of the BID recruitment and participation made the identification (and therefore targeted reminder) of individual participants (who had not yet participated in the second round) impossible, despite emailing general follow-up participation reminders. Given these circumstances, we consider our sample size acceptable, especially in view of the fact that participants had to perform the task twice. Finally, the specific hypotheses were based on the findings of the more adequately powered study 1, and were supported by these preliminarily analyses. Nevertheless, the number of participants and the statistical power were rather low, and a replication study in a larger sample size would prove informative.

The focus of the present studies was on the interaction between the manipulation of sensorimotor states and LAMP. However, other factors such as (upper or lower) limb, laterality, constistency of the laterality of the stimuli with the side of the (desired) amputation, solidity, and perspective, were not considered, given the small number of participants. For the interested reader, we thus provide a table with the results of the explorative analyses including all these factors in the supplementary materials and, for further analyses, the datasets are deposited on the open science framework (OSF, https://osf.io/4z6yc/).

To conclude, our findings extend the accumulating evidence for the functional role of sensorimotor processing in perception, by showing a systematic influence of the sensorimotor state on LAMP. They are thus in line with the embodied cognition framework, suggesting a strong anchoring of perception and cognition in sensorimotor bodily states.

## Acknowledgment

GS received funding from the Swiss National Science Foundation (PP00P1_170511, and P1ZHP1_181383). JTH and BL were funded by funded by the Swiss National Science Foundation (PP00P1_170511). RBB received funding from the Deutsche Forschungsgemeinschaft (DFG; BE 5723/4-1). We also thank Dr. Yannick Rothacher for the useful statistical advices.

