## Supplementary materials for "Limb apparent motion perception: modification by tDCS, and clinically or experimentally altered bodily states"

### Study 1

Table 1. Mean and standard deviation for the combination of the levels of the factors tDCS and SOA

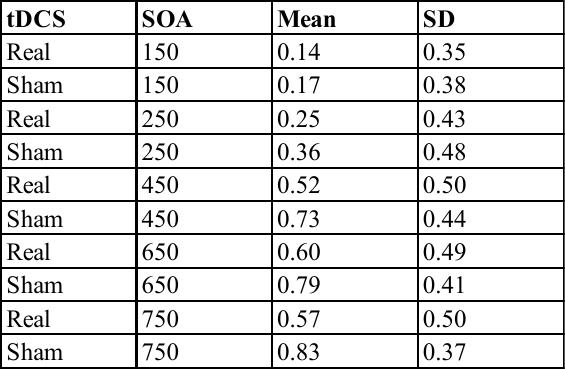

#### Supplementary analyses

i) A linear mixed model inspected the illusion experience as a function of the factors tDCS stimulation (experimental/sham) and SOA (150, 250, 450, 650, 750) and laterality (left/right). The models included the main effects and all the possible two-way and three way interactions. Results are reported in table 2 supplementary materials

Table 2. Results of the linear mixed model examining the impact on the illusion experience in the LAMP task of tDCS stimulation, SOA, and laterality.

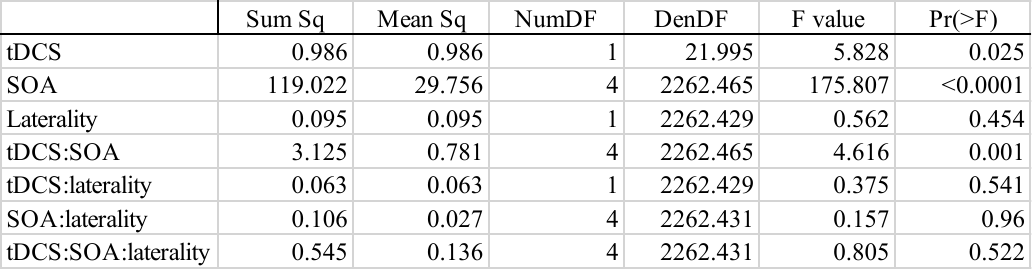

### Study 2

Separately for each of the three samples, we performed an explorative analysis consisting in a linear mixed model where the illusion experience was inspected as a function of the factors sensorimotor state (in individuals with an amputation: wearing the prosthesis/not wearing the prosthesis; individuals with BID: sitting normally/pretending; able-bodied participants: sitting normally/sitting on one leg) , SOA (150, 250, 450, 650, 750), limb (upper/lower limbs), consistency of the affected/immobilised leg with the laterality of the stimuli (“consistency amputation”, consistent/non consistent;), solidity (object/limb solidity), perspective (1pp/3pp). The models included the main effects and all the possible interactions between these factors.

Results for individuals with an amputation, individuals with BID, and able-bodied individuals are reported in table 3, table 4, and table 5 respectively.

Table 3 Results of the linear mixed model examining the impact on the illusion experience in the LAMP task in individuals with a lower limb amputation of sensorimotor state, SOA, limb, consistency amputation, solidity, and perspective

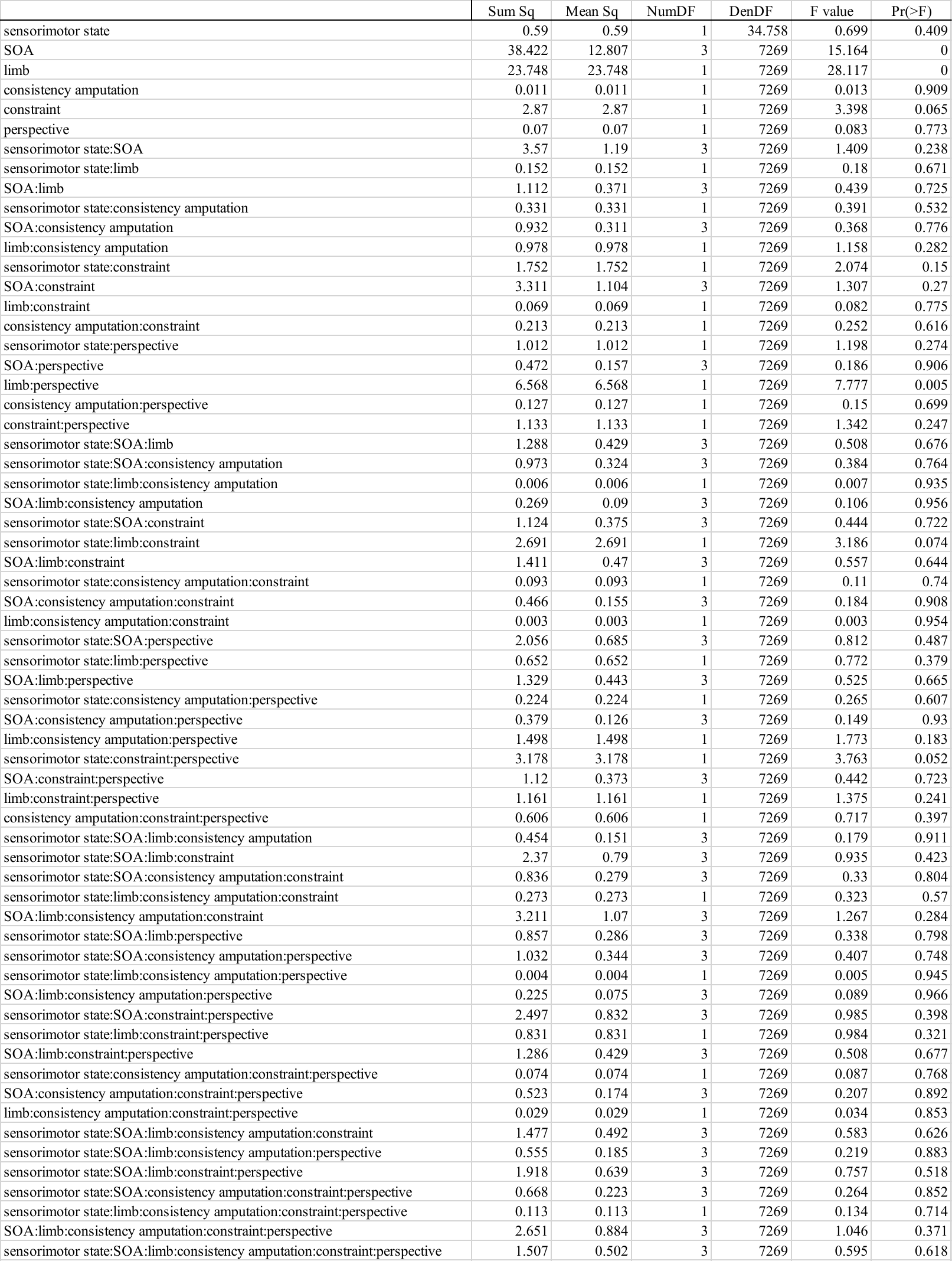

Table 4. Results of the linear mixed model examining the impact on the illusion experience in the LAMP task in individuals with BIID of sensorimotor state, SOA, limb, consistency amputation, solidity, and perspective.

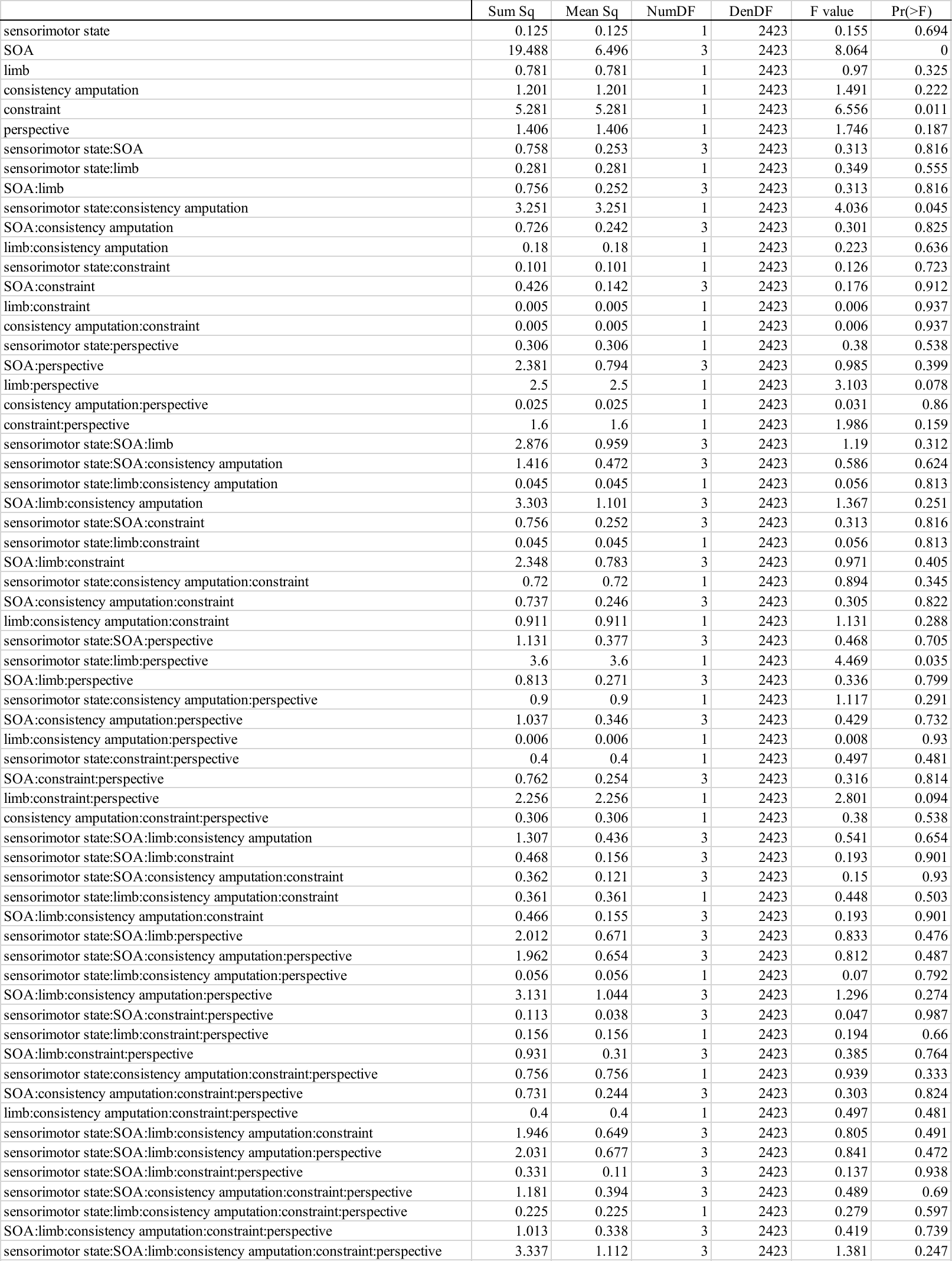

Table 5. Results of the linear mixed model examining the impact on the illusion experience in the LAMP task in able-bodied participants of sensorimotor state, SOA, limb, consistency amputation, solidity, and perspective.

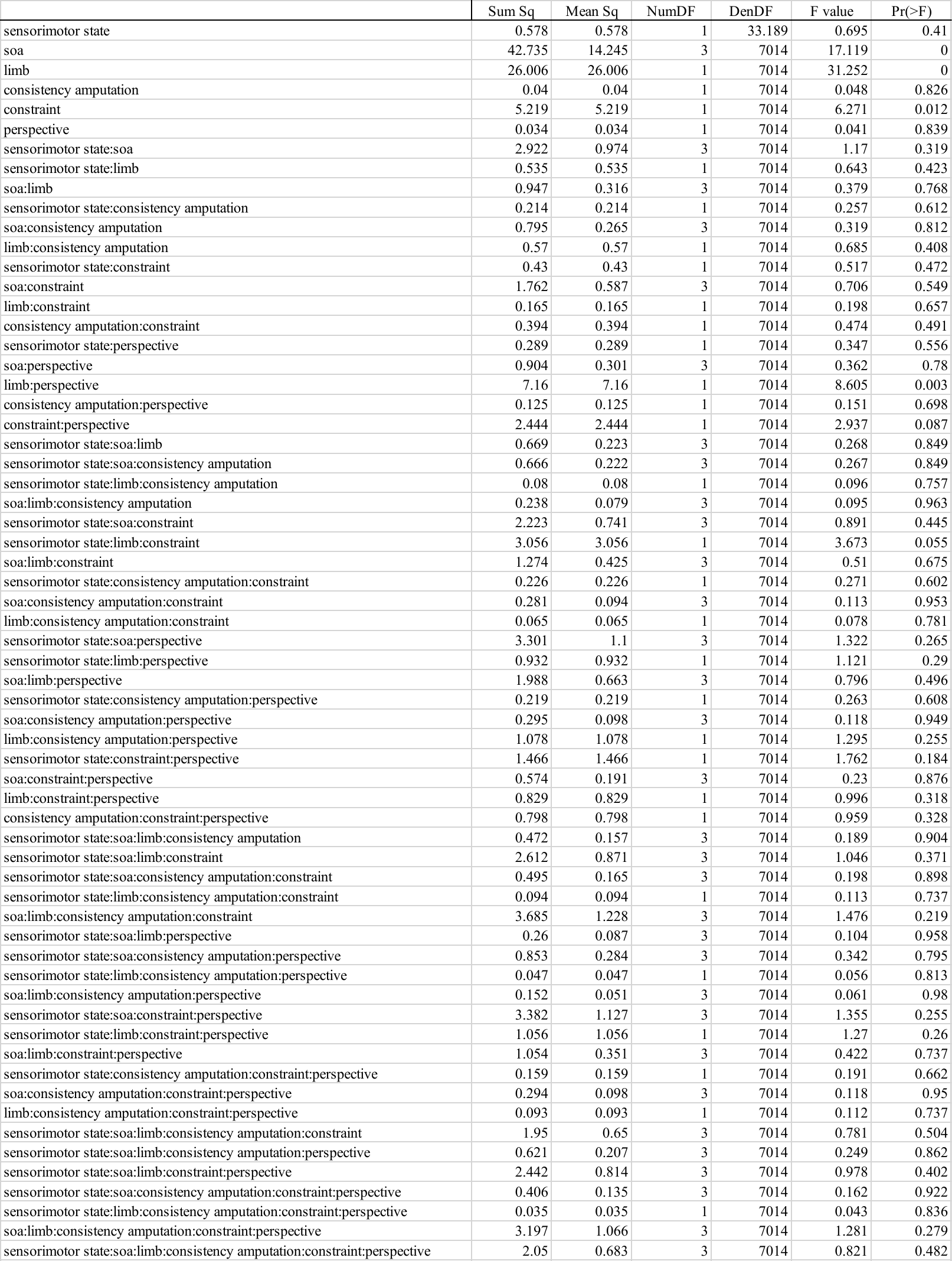
